## Supplemental table 1 to 3 for "Distinct T Cell Receptor (TCR) gene segment usage and MHC-restriction between foetal and adult thymus"

Supplementary Tables

**Supplementary Table 1 Antibody panel for staining thymus from 4 week mice.**

| Antibody | Clone | Supplier | Catalogue number |
|---|---|---|---|
| PE anti-mouse CD3 | 17AD | BioLegend | 100206 |
| PerCP/Cyanine5.5 anti-mouse CD4 | RM4-4 | BioLegend | 116012 |
| FITC anti-mouse CD8a | 53-6.7 | BioLegend | 100706 |

**Supplementary Table 2 Antibody panel for staining thymus from E18.5.**

| Antibody | Clone | Supplier | Catalogue number |
|---|---|---|---|
| Brilliant Violet 421 anti-mouse CD3 | 145-2C11 | BioLegend | 100335 |
| APC anti-mouse CD4 | RM4-4 | BioLegend | 116014 |
| FITC anti-mouse CD8a | 53-6.7 | BioLegend | 100706 |
| PE anti-mouse CD69 | H1.2F3 | eBioscience | 12-0691-83 |

**Supplementary Table 3 | Primers used for TCR sequencing protocol**

| Name | Sequence | Description | Purification |
|---|---|---|---|
| TRAC3 | GAGACCGAGGATCTTTTAACTGG | RT | desalted |
| TRBC2 | GCTTTTGATGGCTCAAACAAGG | RT | desalted |
| 6N_I8.1_6N_M13_2 | [Phos]NNNNNNATCACGACNNNNNNCCAGGGTTTTCCCAGTCACGAC [SpcC3] | Ligation | HPLC |
| malphaRC1 | CAGCAGGTTCTGGGTTCTGGATG | PCR1 | desalted |
| mbetaRC1 | GGGTGGAGTCACATTTCTCAGATCC | PCR1 | desalted |
| SP2_M13 | TTC AGA CGT GTG CTC TTC CGA TCT GTC GTG ACT GGG AAAA CCC TGG | PCR1 | desalted |
| P5-SP1 | AATGATACGGCGACCACCGAGATCTACACTCTTTCCCTACACGACGCTCTTCC | PCR2 | desalted |
| mSP1-6N-I-6-aRC1 | ACACTCTTTCCCTACACGACGCTCTTCCGATCTNNNNNNGCCAATCAGCAGGTTCTGGGTTCTGGATG | SP1 | desalted |
| mSP1-6N-I-6-bRC1 | ACACTCTTTCCCTACACGACGCTCTTCCGATCTNNNNNNGCCAATGGGTGGAGTCACATTTCTCAGATCC | SP1 | desalted |
| mSP1-6N-I-7-aRC1 | ACACTCTTTCCCTACACGACGCTCTTCCGATCTNNNNNNCAGATCCAGCAGGTTCTGGGTTCTGGATG | SP1 | desalted |
| mSP1-6N-I-7-bRC1 | ACACTCTTTCCCTACACGACGCTCTTCCGATCTNNNNNNCAGATCGGGTGGAGTCACATTTCTCAGATCC | SP1 | desalted |
| mSP1-6N-I-8-aRC1 | ACACTCTTTCCCTACACGACGCTCTTCCGATCTNNNNNNACTTGACAGCAGGTTCTGGGTTCTGGATG | SP1 | desalted |
| mSP1-6N-I-8-bRC1 | ACACTCTTTCCCTACACGACGCTCTTCCGATCTNNNNNNACTTGA GGGTGGAGTCACATTTCTCAGATCC | SP1 | desalted |
| mSP1-6HN-I1-aRC1 | ACACTCTTTCCCTACACGACGCTCTTCCGATCTHNHNNHATCACGCAGCAGGTTCTGGGTTCTGGATG | SP1 | desalted |
| mSP1-6HN-I1-bRC1 | ACACTCTTTCCCTACACGACGCTCTTCCGATCTHNHNNHATCACGGGGTGGAGTCACATTTCTCAGATCC | SP1 | desalted |
| mSP1-6HN-I2-aRC1 | ACACTCTTTCCCTACACGACGCTCTTCCGATCTHNHNNHCGATGTCAGCAGGTTCTGGGTTCTGGATG | SP1 | desalted |
| mSP1-6HN-I2-bRC1 | ACACTCTTTCCCTACACGACGCTCTTCCGATCTHNHNNHCGATGTGGGTGGAGTCACATTTCTCAGATCC | SP1 | desalted |
| mSP1-6HN-I3-aRC1 | ACACTCTTTCCCTACACGACGCTCTTCCGATCTHNHNNHTTAGGCCAGCAGGTTCTGGGTTCTGGATG | SP1 | desalted |
| mSP1-6HN-I3-bRC1 | ACACTCTTTCCCTACACGACGCTCTTCCGATCTHNHNNHTTAGGCGGGTGGAGTCACATTTCTCAGATCC | SP1 | desalted |
| mSP1-6HN-I4-aRC1 | ACACTCTTTCCCTACACGACGCTCTTCCGATCTHNHNNHTGACCACAGCAGGTTCTGGGTTCTGGATG | SP1 | desalted |
| mSP1-6HN-I4-bRC1 | ACACTCTTTCCCTACACGACGCTCTTCCGATCTHNHNNHTGACCAGGGTGGAGTCACATTTCTCAGATCC | SP1 | desalted |
| mSP1-6HN-I5-aRC1 | ACACTCTTTCCCTACACGACGCTCTTCCGATCTHNHNNHACAGTGCAGCAGGTTCTGGGTTCTGGATG | SP1 | desalted |
| mSP1-6HN-I5-bRC1 | ACACTCTTTCCCTACACGACGCTCTTCCGATCTHNHNNHACAGTGGGGTGGAGTCACATTTCTCAGATCC | SP1 | desalted |
| mSP1-6HN-I6-aRC1 | ACACTCTTTCCCTACACGACGCTCTTCCGATCTHNHNNHGCCAATCAGCAGGTTCTGGGTTCTGGATG | SP1 | desalted |

| | | | |
|---|---|---|---|
| mSP1-6HN-I6-bRC1 | ACACTCTTTCCCTACACGACGCTCTTCCGATCTHNHNNHGCCAATGGGTGGAGTCACATTTCTCAGATCC | SP1 | desalted |
| mSP1-6HN-I7-aRC1 | ACACTCTTTCCCTACACGACGCTCTTCCGATCTHNHNNHCAGATCCAGCAGGTTCTGGGTTCTGGATG | SP1 | desalted |
| mSP1-6HN-I7-bRC1 | ACACTCTTTCCCTACACGACGCTCTTCCGATCTHNHNNHCAGATCGGGTGGAGTCACATTTCTCAGATCC | SP1 | desalted |
| mSP1-6HN-I8-aRC1 | ACACTCTTTCCCTACACGACGCTCTTCCGATCTHNHNNHACTTGACAGCAGGTTCTGGGTTCTGGATG | SP1 | desalted |
| mSP1-6HN-I8-bRC1 | ACACTCTTTCCCTACACGACGCTCTTCCGATCTHNHNNHACTTGAGGGTGGAGTCACATTTCTCAGATCC | SP1 | desalted |
| mSP1-6HN-I9-aRC1 | ACACTCTTTCCCTACACGACGCTCTTCCGATCTHNHNNHGATCAGCAGCAGGTTCTGGGTTCTGGATG | SP1 | desalted |
| mSP1-6HN-I9-bRC1 | ACACTCTTTCCCTACACGACGCTCTTCCGATCTHNHNNHGATCAGGGGTGGAGTCACATTTCTCAGATCC | SP1 | desalted |
| mSP1-6HN-I10-aRC1 | ACACTCTTTCCCTACACGACGCTCTTCCGATCTHNHNNHTAGCTTCAGCAGGTTCTGGGTTCTGGATG | SP1 | desalted |
| mSP1-6HN-I10-bRC1 | ACACTCTTTCCCTACACGACGCTCTTCCGATCTHNHNNHTAGCTTGGGTGGAGTCACATTTCTCAGATCC | SP1 | desalted |
| mSP1-6HN-I11-aRC1 | ACACTCTTTCCCTACACGACGCTCTTCCGATCTHNHNNHGGCTACCAGCAGGTTCTGGGTTCTGGATG | SP1 | desalted |
| mSP1-6HN-I11-bRC1 | ACACTCTTTCCCTACACGACGCTCTTCCGATCTHNHNNHGGCTACGGGTGGAGTCACATTTCTCAGATCC | SP1 | desalted |
| mSP1-6HN-I12-aRC1 | ACACTCTTTCCCTACACGACGCTCTTCCGATCTHNHNNHCTTGTACAGCAGGTTCTGGGTTCTGGATG | SP1 | desalted |
| mSP1-6HN-I12-bRC1 | ACACTCTTTCCCTACACGACGCTCTTCCGATCTHNHNNHCTTGTAGGGTGGAGTCACATTTCTCAGATCC | SP1 | desalted |
| mSP1-6HN-I13-aRC1 | ACACTCTTTCCCTACACGACGCTCTTCCGATCTHNHNNHTAGACTCAGCAGGTTCTGGGTTCTGGATG | SP1 | desalted |
| mSP1-6HN-I13-bRC1 | ACACTCTTTCCCTACACGACGCTCTTCCGATCTHNHNNHTAGACTGGGTGGAGTCACATTTCTCAGATCC | SP1 | desalted |
| P7-I8.15_SP2 | CAAGCAGAAGACGGCATACGAGATCTACCAGGGTGACTGGAGTTCAGACGTGTGCTCTTCCGATC | SP2 P7 | desalted |
| P7-I8.16_SP2 | CAAGCAGAAGACGGCATACGAGATCATGCTTAGTGACTGGAGTTCAGACGTGTGCTCTTCCGATC | SP2 P7 | desalted |
| P7-I8.17_SP2 | CAAGCAGAAGACGGCATACGAGATGCACATCTGTGACTGGAGTTCAGACGTGTGCTCTTCCGATC | SP2 P7 | desalted |
| P7-I8.18_SP2 | CAAGCAGAAGACGGCATACGAGATTGCTCGACGTGACTGGAGTTCAGACGTGTGCTCTTCCGATC | SP2 P7 | desalted |
| P7-I8.19_SP2 | CAAGCAGAAGACGGCATACGAGATAGCAATTCGTGACTGGAGTTCAGACGTGTGCTCTTCCGATC | SP2 P7 | desalted |
| P7-I8.20_SP2 | CAAGCAGAAGACGGCATACGAGATAGTTGCTTGTGACTGGAGTTCAGACGTGTGCTCTTCCGATC | SP2 P7 | desalted |
| P7-I8.21_SP2 | CAAGCAGAAGACGGCATACGAGATCCAGTTAGGTGACTGGAGTTCAGACGTGTGCTCTTCCGATC | SP2 P7 | desalted |
| P7-I8.22_SP2 | CAAGCAGAAGACGGCATACGAGATTTGAGCCTGTGACTGGAGTTCAGACGTGTGCTCTTCCGATC | SP2 P7 | desalted |
| P7-I8.23_SP2 | CAAGCAGAAGACGGCATACGAGATACCAACTGGTGACTGGAGTTCAGACGTGTGCTCTTCCGATC | SP2 P7 | desalted |
| P7-I8.24_SP2 | CAAGCAGAAGACGGCATACGAGATGGTCCAGAGTGACTGGAGTTCAGACGTGTGCTCTTCCGATC | SP2 P7 | desalted |

| | | | |
|---|---|---|---|
| P7-I8.25_SP2 | CAAGCAGAAGACGGCATACGAGATGTATAACAGTGACTGGAGTTCAGACGTGTGCTCTTCCGATC | SP2 P7 | desalted |
| P7-I8.26_SP2 | CAAGCAGAAGACGGCATACGAGATTTCGCTGAGTGACTGGAGTTCAGACGTGTGCTCTTCCGATC | SP2 P7 | desalted |
| P7_L27 | CAAGCAGAAGACGGCATACGAGAT CGCTATGT GTGACTGGAGTTCAGACGTGTGCTCTTCCGATC | SP2 P7 | desalted |
| P7_L28 | CAAGCAGAAGACGGCATACGAGAT TAAGCACA GTGACTGGAGTTCAGACGTGTGCTCTTCCGATC | SP2 P7 | desalted |
| P7_L29 | CAAGCAGAAGACGGCATACGAGAT GTAACATC GTGACTGGAGTTCAGACGTGTGCTCTTCCGATC | SP2 P7 | desalted |
| P7_L30 | CAAGCAGAAGACGGCATACGAGAT ACTAAGAC GTGACTGGAGTTCAGACGTGTGCTCTTCCGATC | SP2 P7 | desalted |
| P7_L31 | CAAGCAGAAGACGGCATACGAGAT TGTAACTC GTGACTGGAGTTCAGACGTGTGCTCTTCCGATC | SP2 P7 | desalted |
| P7_L32 | CAAGCAGAAGACGGCATACGAGAT AACAATGG GTGACTGGAGTTCAGACGTGTGCTCTTCCGATC | SP2 P7 | desalted |
| P7_L33 | CAAGCAGAAGACGGCATACGAGAT CCTTCGCA GTGACTGGAGTTCAGACGTGTGCTCTTCCGATC | SP2 P7 | desalted |
| P7_L34 | CAAGCAGAAGACGGCATACGAGAT GACCGTTG GTGACTGGAGTTCAGACGTGTGCTCTTCCGATC | SP2 P7 | desalted |
| P7_L35 | CAAGCAGAAGACGGCATACGAGAT TCTGCAAG GTGACTGGAGTTCAGACGTGTGCTCTTCCGATC | SP2 P7 | desalted |
| P7_L36 | CAAGCAGAAGACGGCATACGAGAT CACATCCT GTGACTGGAGTTCAGACGTGTGCTCTTCCGATC | SP2 P7 | desalted |
| P7_L37 | CAAGCAGAAGACGGCATACGAGAT AGGATCTA GTGACTGGAGTTCAGACGTGTGCTCTTCCGATC | SP2 P7 | desalted |
| P7_L38 | CAAGCAGAAGACGGCATACGAGAT GTCATCTA GTGACTGGAGTTCAGACGTGTGCTCTTCCGATC | SP2 P7 | desalted |
| P7_L39 | CAAGCAGAAGACGGCATACGAGAT GAACCTAG GTGACTGGAGTTCAGACGTGTGCTCTTCCGATC | SP2 P7 | desalted |
| P7_L40 | CAAGCAGAAGACGGCATACGAGAT TTACGCAC GTGACTGGAGTTCAGACGTGTGCTCTTCCGATC | SP2 P7 | desalted |
| P7_L41 | CAAGCAGAAGACGGCATACGAGAT AGGTGCGA GTGACTGGAGTTCAGACGTGTGCTCTTCCGATC | SP2 P7 | desalted |
| P7_L42 | CAAGCAGAAGACGGCATACGAGAT CATGATCG GTGACTGGAGTTCAGACGTGTGCTCTTCCGATC | SP2 P7 | desalted |
| P7_L43 | CAAGCAGAAGACGGCATACGAGAT GCCGCAAC GTGACTGGAGTTCAGACGTGTGCTCTTCCGATC | SP2 P7 | desalted |
| P7_L44 | CAAGCAGAAGACGGCATACGAGAT TTATATCT GTGACTGGAGTTCAGACGTGTGCTCTTCCGATC | SP2 P7 | desalted |
| P7_L45 | CAAGCAGAAGACGGCATACGAGAT CTGTGGCG GTGACTGGAGTTCAGACGTGTGCTCTTCCGATC | SP2 P7 | desalted |
| P7_L46 | CAAGCAGAAGACGGCATACGAGAT AACGCATT GTGACTGGAGTTCAGACGTGTGCTCTTCCGATC | SP2 P7 | desalted |
| P7_L47 | CAAGCAGAAGACGGCATACGAGAT AACTTGAC GTGACTGGAGTTCAGACGTGTGCTCTTCCGATC | SP2 P7 | desalted |
| P7_L48 | CAAGCAGAAGACGGCATACGAGAT CGCCTTCC GTGACTGGAGTTCAGACGTGTGCTCTTCCGATC | SP2 P7 | desalted |
| P7_L49 | CAAGCAGAAGACGGCATACGAGAT GACCAGGA GTGACTGGAGTTCAGACGTGTGCTCTTCCGATC | SP2 P7 | desalted |

| | | | |
|---|---|---|---|
| P7_L50 | CAAGCAGAAGACGGCATACGAGAT TCCTTGGT GTGACTGGAGTTCAGACGTGTGCTCTTCCGATC | SP2 P7 | desalted |
| P7_L51 | CAAGCAGAAGACGGCATACGAGAT AAGACACT GTGACTGGAGTTCAGACGTGTGCTCTTCCGATC | SP2 P7 | desalted |
| P7_L52 | CAAGCAGAAGACGGCATACGAGAT CTGTAATC GTGACTGGAGTTCAGACGTGTGCTCTTCCGATC | SP2 P7 | desalted |
| P7_L53 | CAAGCAGAAGACGGCATACGAGAT GAAGAAGT GTGACTGGAGTTCAGACGTGTGCTCTTCCGATC | SP2 P7 | desalted |
| P7_L54 | CAAGCAGAAGACGGCATACGAGAT TAATGAAC GTGACTGGAGTTCAGACGTGTGCTCTTCCGATC | SP2 P7 | desalted |
| P7_L55 | CAAGCAGAAGACGGCATACGAGAT TCCAGCAA GTGACTGGAGTTCAGACGTGTGCTCTTCCGATC | SP2 P7 | desalted |
| P7_L56 | CAAGCAGAAGACGGCATACGAGAT GTCCACAG GTGACTGGAGTTCAGACGTGTGCTCTTCCGATC | SP2 P7 | desalted |
| P7_L57 | CAAGCAGAAGACGGCATACGAGAT CAATAGTC GTGACTGGAGTTCAGACGTGTGCTCTTCCGATC | SP2 P7 | desalted |
| P7_L58 | CAAGCAGAAGACGGCATACGAGAT AGGTAAGG GTGACTGGAGTTCAGACGTGTGCTCTTCCGATC | SP2 P7 | desalted |
| P7_L59 | CAAGCAGAAGACGGCATACGAGAT TACTTAGC GTGACTGGAGTTCAGACGTGTGCTCTTCCGATC | SP2 P7 | desalted |
| P7_L60 | CAAGCAGAAGACGGCATACGAGAT GAAGGAAG GTGACTGGAGTTCAGACGTGTGCTCTTCCGATC | SP2 P7 | desalted |
| P7_L61 | CAAGCAGAAGACGGCATACGAGAT CATAGCGA GTGACTGGAGTTCAGACGTGTGCTCTTCCGATC | SP2 P7 | desalted |
| P7_L62 | CAAGCAGAAGACGGCATACGAGAT ATTGTCTG GTGACTGGAGTTCAGACGTGTGCTCTTCCGATC | SP2 P7 | desalted |
| P7_L63 | CAAGCAGAAGACGGCATACGAGAT CAACTCTC GTGACTGGAGTTCAGACGTGTGCTCTTCCGATC | SP2 P7 | desalted |
| P7_L64 | CAAGCAGAAGACGGCATACGAGAT ATTCTAGG GTGACTGGAGTTCAGACGTGTGCTCTTCCGATC | SP2 P7 | desalted |
| P7_L65 | CAAGCAGAAGACGGCATACGAGAT TGCTGCTG GTGACTGGAGTTCAGACGTGTGCTCTTCCGATC | SP2 P7 | desalted |
| P7_L66 | CAAGCAGAAGACGGCATACGAGAT GCCTAGCC GTGACTGGAGTTCAGACGTGTGCTCTTCCGATC | SP2 P7 | desalted |
| P7_L67 | CAAGCAGAAGACGGCATACGAGAT CCTATGCC GTGACTGGAGTTCAGACGTGTGCTCTTCCGATC | SP2 P7 | desalted |
| P7_L68 | CAAGCAGAAGACGGCATACGAGAT ATAGCGTC GTGACTGGAGTTCAGACGTGTGCTCTTCCGATC | SP2 P7 | desalted |
| P7_L69 | CAAGCAGAAGACGGCATACGAGAT TGTCGGAT GTGACTGGAGTTCAGACGTGTGCTCTTCCGATC | SP2 P7 | desalted |
| P7_L70 | CAAGCAGAAGACGGCATACGAGAT GACAGTAA GTGACTGGAGTTCAGACGTGTGCTCTTCCGATC | SP2 P7 | desalted |
| P7_L71 | CAAGCAGAAGACGGCATACGAGAT GCCGTCGA GTGACTGGAGTTCAGACGTGTGCTCTTCCGATC | SP2 P7 | desalted |
| P7_L72 | CAAGCAGAAGACGGCATACGAGAT TATCCAGG GTGACTGGAGTTCAGACGTGTGCTCTTCCGATC | SP2 P7 | desalted |
| P7_L73 | CAAGCAGAAGACGGCATACGAGAT ATTATGTT GTGACTGGAGTTCAGACGTGTGCTCTTCCGATC | SP2 P7 | desalted |
| P7_L74 | CAAGCAGAAGACGGCATACGAGAT CCAACATT GTGACTGGAGTTCAGACGTGTGCTCTTCCGATC | SP2 P7 | desalted |

| | | | |
|---|---|---|---|
| P7_L75 | CAAGCAGAAGACGGCATACGAGAT GATATCCA GTGACTGGAGTTCAGACGTGTGCTCTTCCGATC | SP2 P7 | desalted |
| P7_L76 | CAAGCAGAAGACGGCATACGAGAT TGCAAGTA GTGACTGGAGTTCAGACGTGTGCTCTTCCGATC | SP2 P7 | desalted |
| P7_L77 | CAAGCAGAAGACGGCATACGAGAT AATGTTCT GTGACTGGAGTTCAGACGTGTGCTCTTCCGATC | SP2 P7 | desalted |
| P7_L78 | CAAGCAGAAGACGGCATACGAGAT CAGCGGTA GTGACTGGAGTTCAGACGTGTGCTCTTCCGATC | SP2 P7 | desalted |
| P7_L79 | CAAGCAGAAGACGGCATACGAGAT ATTCCTCT GTGACTGGAGTTCAGACGTGTGCTCTTCCGATC | SP2 P7 | desalted |
| P7_L80 | CAAGCAGAAGACGGCATACGAGAT CTGCGGAT GTGACTGGAGTTCAGACGTGTGCTCTTCCGATC | SP2 P7 | desalted |
| P7_L81 | CAAGCAGAAGACGGCATACGAGAT GTCTGATG GTGACTGGAGTTCAGACGTGTGCTCTTCCGATC | SP2 P7 | desalted |
| P7_L82 | CAAGCAGAAGACGGCATACGAGAT TATCTGCC GTGACTGGAGTTCAGACGTGTGCTCTTCCGATC | SP2 P7 | desalted |
| P7_L83 | CAAGCAGAAGACGGCATACGAGAT ACACGATC GTGACTGGAGTTCAGACGTGTGCTCTTCCGATC | SP2 P7 | desalted |
| P7_L84 | CAAGCAGAAGACGGCATACGAGAT CCAGAGCT GTGACTGGAGTTCAGACGTGTGCTCTTCCGATC | SP2 P7 | desalted |
| P7_L85 | CAAGCAGAAGACGGCATACGAGAT GACCTAAC GTGACTGGAGTTCAGACGTGTGCTCTTCCGATC | SP2 P7 | desalted |
| P7_L86 | CAAGCAGAAGACGGCATACGAGAT TCGCCTTG GTGACTGGAGTTCAGACGTGTGCTCTTCCGATC | SP2 P7 | desalted |
| P7_L87 | CAAGCAGAAGACGGCATACGAGAT CCTACCAT GTGACTGGAGTTCAGACGTGTGCTCTTCCGATC | SP2 P7 | desalted |
| P7_L88 | CAAGCAGAAGACGGCATACGAGAT TCGCTAGA GTGACTGGAGTTCAGACGTGTGCTCTTCCGATC | SP2 P7 | desalted |
| P7_L89 | CAAGCAGAAGACGGCATACGAGAT AAGGATGT GTGACTGGAGTTCAGACGTGTGCTCTTCCGATC | SP2 P7 | desalted |
| P7_L90 | CAAGCAGAAGACGGCATACGAGAT CTAACTCG GTGACTGGAGTTCAGACGTGTGCTCTTCCGATC | SP2 P7 | desalted |
| P7_L91 | CAAGCAGAAGACGGCATACGAGAT ACAGGTAT GTGACTGGAGTTCAGACGTGTGCTCTTCCGATC | SP2 P7 | desalted |
| P7_L92 | CAAGCAGAAGACGGCATACGAGAT TCTCGGTC GTGACTGGAGTTCAGACGTGTGCTCTTCCGATC | SP2 P7 | desalted |
| P7_L93 | CAAGCAGAAGACGGCATACGAGAT ACAGTTGA GTGACTGGAGTTCAGACGTGTGCTCTTCCGATC | SP2 P7 | desalted |
| P7_L94 | CAAGCAGAAGACGGCATACGAGAT CTATGCGT GTGACTGGAGTTCAGACGTGTGCTCTTCCGATC | SP2 P7 | desalted |
| P7_L95 | CAAGCAGAAGACGGCATACGAGAT CAGGAGCC GTGACTGGAGTTCAGACGTGTGCTCTTCCGATC | SP2 P7 | desalted |
| P7_L96 | CAAGCAGAAGACGGCATACGAGAT AGGTCGCA GTGACTGGAGTTCAGACGTGTGCTCTTCCGATC | SP2 P7 | desalted |
| P7_L97 | CAAGCAGAAGACGGCATACGAGAT CAGCAAGG GTGACTGGAGTTCAGACGTGTGCTCTTCCGATC | SP2 P7 | desalted |
| P7_L98 | CAAGCAGAAGACGGCATACGAGAT ATTATCAA GTGACTGGAGTTCAGACGTGTGCTCTTCCGATC | SP2 P7 | desalted |
| P7_L99 | CAAGCAGAAGACGGCATACGAGAT TTAATCAG GTGACTGGAGTTCAGACGTGTGCTCTTCCGATC | SP2 P7 | desalted |

| | | | |
|---|---|---|---|
| P7_L100 | CAAGCAGAAGACGGCATACGAGAT CGTTACCA GTGACTGGAGTTCAGACGTGTGCTCTTCCGATC | SP2 P7 | desalted |
| P7_L101 | CAAGCAGAAGACGGCATACGAGAT AAGTAGAG GTGACTGGAGTTCAGACGTGTGCTCTTCCGATC | SP2 P7 | desalted |
| P7_L102 | CAAGCAGAAGACGGCATACGAGAT TTGAATAG GTGACTGGAGTTCAGACGTGTGCTCTTCCGATC | SP2 P7 | desalted |
| P5 | AATGATACGGCGACCACCGAGATCTACACT | QPCR | desalted |
| P7 | CAAGCAGAAGACGGCATACGAGAT | QPCR | desalted |

**Supplementary Table 4 | Reagents and consumables for TCR sequencing protocol**

| Reagent | Supplier | Cat. No. |
|---|---|---|
| RQ1 RNase-Free DNase | Promega | M6101 |
| RQ1 DNase 10X Reaction Buffer | Promega | M6101 |
| RQ1 Dnase Stop Solution | Promega | M6101 |
| RNase free water | Invitrogen | 10977-035 |
| dNTPs (10 mM) | Promega | U1515 |
| SuperScript III RT (200 U/µl) | Invitrogen | 18080085 |
| 5X First-Strand (FS) Buffer | Invitrogen | 18080085 |
| 0.1 M DTT | Invitrogen | 18080085 |
| RNasin | Promega | N2115 |
| Minelute PCR Purification kit | Qiagen | 28006 |
| T4 RNA Ligase Reaction Buffer | NEB | M0204L |
| Adenosine-5'-Triphosphate (ATP) | NEB | M0204L |
| PEG 8000 | NEB | M0204L |
| T4 RNA Ligase 1 (ssRNA Ligase) | NEB | M0204L |
| BSA (20 mg/mL) | NEB | B9000S |
| Hexammine cobalt(III) chloride (HCC) | Sigma | H7891-5G |
| Agencourt AMPure beads XP | Beckman Coulter | A63881 |
| 5x Phusion HF buffer | NEB | M0530L |
| Phusion Polymerase | NEB | M0530L |
| ROX Reference Dye | Invitrogen | 12223012 |
| SYBR Green I Nucleic Acid Gel Stain 10,000× | Invitrogen | S7563 |
| DMSO for molecular biology | Sigma | D8418-50ML |
| Qubit dsDNA HS Assay Kit | ThermoFisher Scientific | Q32854 |
| Qubit Assay Tubes | ThermoFisher Scientific | Q32856 |
| High Sensitivity D1000 ScreenTape | Agilent | 5067-5584 |
| High Sensitivity D1000 Reagents | Agilent | 5067-5585 |
| High Sensitivity D1000 Ladder | Agilent | 5067-5587 |
| 96-well Plates | Agilent | 5042-8502 |
| 96-well Plate Foil Seal | Agilent | 5067-5154 |
| Pippin Gel Cassette 1.5% agarose dye free 250bp-1.5kb | Sage Science | CDF1510 |
| PhiX Control V3 | Illumina | FC-110-3001 |
| MiSeq Reagent Kit v2 (500-cycles) | Illumina | MS-102-2003 |
| Eppendorf DNA LoBind Polypropylene Microcentrifuge Tube | FisherScientific | 10051232 |
| 96-well PCR plate semi skirted | Starlab | I1402-9700C |
| Adhesive PCR Plate Seal | SLS | 4ti-0500 |
| PicoPure RNA Isolation Kit | Applied Biosystems | KIT0204 |
| 500/550 Mid Output Kit v2.5 (300 Cycles) | Illumina | 20024905 |
| Blitz Away RNase spray | Serem Biotech | 40-1735-10 |
| Tris (1M), pH 7.0, RNase-free | ThermoFisher Scientific | AM9850G |

| | | |
|---|---|---|
| Sodium hydroxide solution for molecular biology, 10M in H2O | Sigma | 72068-100ML |
